# High throughput chromatographic ultra-purification of virus-like particles for downstream viromics

**DOI:** 10.64898/2026.07.09.737491

**Authors:** Jessie Maier, Namita Deshmukh, Manuel Kleiner

## Abstract

Virus-like particles (VLPs) are an abundant component of microbiomes with critical ecological roles such as population control through viral predation and horizontal gene transfer. Studying the collection of viruses in microbiomes (the virome) through metagenomics has provided important insights into the composition and functions of VLPs in different environments. However, the current gold-standard method for VLP purification, CsCl density gradient ultracentrifugation (CsCl), is low throughput, time consuming and suffers from biases which limits the ability to study viromes in larger sample sets and can interfere with data interpretation. Here we present an anion exchange (AEX) chromatography-based approach for the purification of VLPs from microbiome samples that allows for significant increases in throughput and reproducibility while achieving VLP purity levels similar to or higher than CsCl. We used microbiome samples of known composition to first establish and evaluate the AEX approaches and compare them to CsCl. We implemented the AEX approach both for fast performance liquid chromatography (FPLC) and in multi-well plates. We compared the VLPs purified with CsCl and AEX using shotgun metagenomic sequencing and found that AEX performs similarly to or better than CsCl for purification of VLPs. AEX purified VLP-fractions captured significantly more viral DNA compared to CsCl. We also found that both AEX and CsCl were capable of capturing viruses present at extremely low relative abundances (<0.001%). Additionally, we found that DNase digestion and CsCl may bias against filamentous phage morphologies. Finally, we purified VLPs from conventional murine feces using AEX and CsCl. AEX purified murine fecal VLPs had a much higher viral DNA content (85%) than CsCl (41%). While there were some differences in viral contigs assembled from AEX and CsCl VLP metagenomes, these method unique viral contigs made up only small proportions (<8%) of the relative abundance in the VLP metagenomes. AEX, particularly in the multi-well format, enables the ultrapurification of VLPs from tens to hundreds of samples in a single day thus facilitating virome studies with the large sample numbers needed for translational and clinical research.

## Introduction

Virus-like particles, VLPs, are defined as extracellular particles resembling viruses that may, or may not be, infectious and carry nucleic acid^1^. This general term was developed to describe filterable particles that fluoresce under epifluorescence microscopy with a DNA stain and therefore encompasses a wide repertoire of both viral and ‘virus-likè particles. Non-viral VLPs include ultrasmall filterable bacteria^2^, gene transfer agents (GTAs)^3^ and extracellular membrane vesicles (MVs)^4^, all of which are non-infectious particles that carry bacterial DNA. In most environments, the majority of VLPs are bacterial viruses (bacteriophage) while eukaryotic viruses and other VLPs make up a smaller proportion^5–7^. Bacteriophage (phage) play critical roles in microbiomes by controlling bacterial populations and contributing to nutrient cycling through predation and lysis^8^, augmenting host fitness through auxiliary metabolic genes and lysogenic conversion^9,10^, and facilitating rapid and dynamic horizontal gene transfer through transduction^11^. Phage are becoming increasingly recognized as a catalyst for altering microbiome composition^12^ and function^13^ and even host physiology^14–16^ in mammalian hosts.

Due to the importance of the collection of viruses, including phages, in microbiomes, more and more scientists are turning towards the study of viromes using metagenomic sequencing. To obtain “clean” metagenomes of viromes that truly represent nucleic acid carried by VLPs rather than contamination from cellular organisms, VLPs must be ultra-purified from the associated microbiome. The current gold-standard method for VLP ultra-purification is cesium chloride density gradient ultra-centrifugation (CsCl), which usually follows an initial concentration, filtering and DNAse digest^17,18^. The use of density gradient ultra-centrifugation for VLP purification represents a major bottleneck for virome studies as only a limited number of samples can be purified in an ultra-centrifuge. Commonly, ultra-centrifuges have space for 6 samples and one purification run takes anywhere between 24-36 hours thus limiting sample throughput enormously^17^. This often limits virome studies to small sample numbers^19–22^ and prevents larger translational or clinical virome studies from using ultra-purified VLPs. Additionally, density gradient ultra-centrifugation has been shown to strongly bias against specific phages^18,23^ which could be due to the fact that usually only one density interface is harvested or sensitivity of some phage groups against CsCl^17^.

In summary, the current methods used to purify VLPs from microbiome samples for metagenomic sequencing are inefficient, low throughput and suffer from biases. Relatively recently, liquid chromatography methods have been developed to efficiently concentrate and purify single VLPs from pure cultures of bacteriophages or eukaryotic viruses^24^. While traditional resin-based columns are not designed for large molecular weight particles like VLPs, monolithic columns, made from a porous material with large channels, are more suited for VLPs. The large channel size on monolithic columns reduces back pressure and associated shear forces and prevents exclusion of large biomolecules^24–26^. Additionally, monolithic columns are flow rate independent which means that purifications can be scaled appropriately without concern for change in resolution. Monolithic chromatography has already been recognized as an alternative to density gradients for purification of single VLPs^27^ and many studies have successfully used monoliths for this purpose^27–31^. Despite the routine use of monoliths for VLP purification from relatively simple sample matrices, the application of monoliths for simultaneous purification of diverse VLPs from complex sample matrices, such as generally required in viromics studies, has not been thoroughly assessed^32^. We performed an evaluation of fast performance liquid chromatography (FPLC) using monolithic anion exchange (AEX) columns versus the gold-standard CsCl density gradient ultra-centrifugation method columns for the ultra-purification of

VLPs from complex sample matrices. For initial method evaluation we used microbiome samples of known composition generated from fecal material of germ-free mice inoculated with a known 14-member microbial community and a spike-in of five bacteriophages. Then we demonstrated the use of the AEX approach using fecal samples from mice with a conventional microbiome. We focused on determining the purity of VLPs in terms of contamination with free (bacterial, mouse) DNA, the number of different viral genomes recovered, and their relative abundance. Since the requirement for an FPLC limits throughput and accessibility for the FPLC-based method, we also tested AEX columns in a 24-well plate format (Well).

## Results and Discussion

### Evaluation and comparison of AEX chromatography methods to CsCl using fecal samples with a defined microbiota and phage spike-ins

We developed a microbiome sample that mimicked the complexity of a fecal sample albeit with known content of bacteria and phage so we could evaluate the AEX-based VLP purification methods as compared to CsCl. We spiked a known phage mix consisting of P22, M13, ES18, T4 and T3 into fecal homogenates generated from pellets collected from gnotobiotic mice gavaged with a 14-member defined bacterial community^33^. The phage chosen represent different morphologies (podoviridae, siphoviridae, myoviridae, and filamentous) and sizes (Table S1). We purified VLPs from these samples with AEX using an FPLC (LC) and CsCl (Fig. 1). We also replicated the AEX LC method with 24-well plates (Well) containing the same AEX column material used in the FPLC purifications. Samples were pre-treated by centrifugation, filtration and DNAse digestion as commonly performed in virome studies (Fig. 1). We used metagenomic shotgun sequencing to characterize the DNA content of samples along the sample processing pipeline (n=4) after the filtration step which has been shown to remove a large share of bacterial DNA through removal of intact cells^18,34,35^.

**Figure 1.**
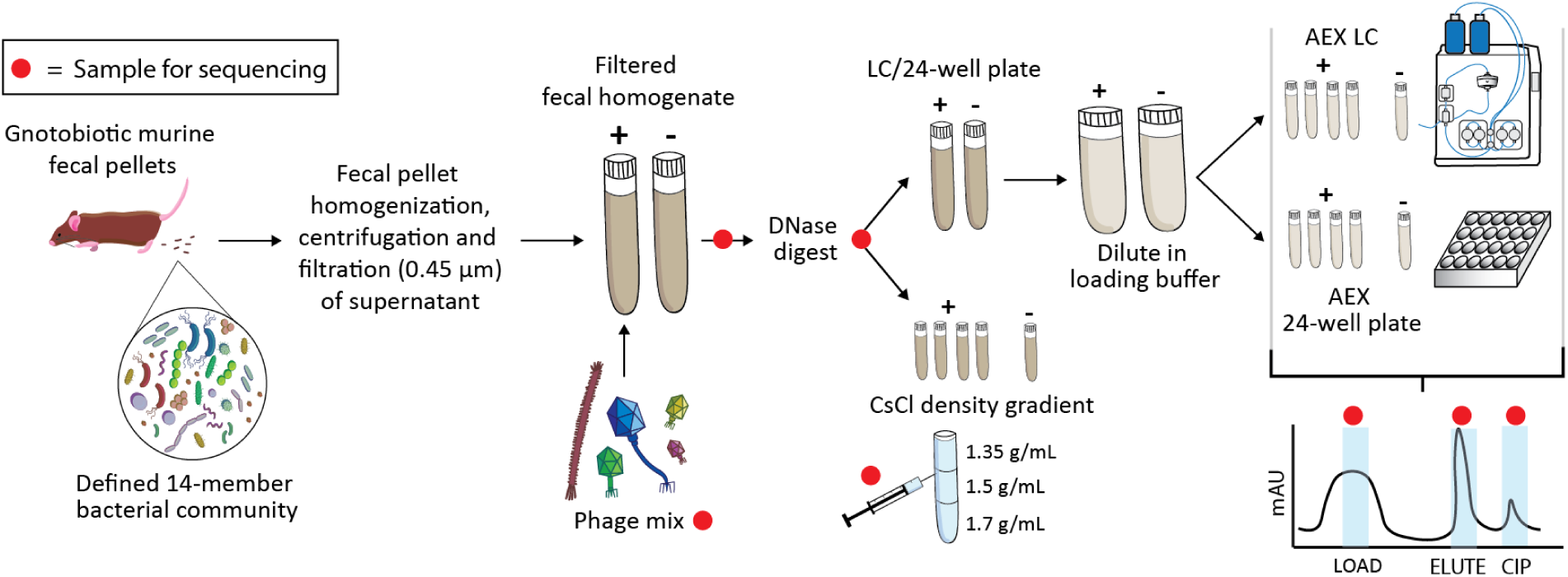
Experimental overview for purification of five phages spiked into fecal homogenates from gnotobiotic mice gavaged with a defined community consisting of 14 bacteria. The phage mix contained phage P22, ES18, M13, T3 and T4. The defined microbiota contained *Akkermanisa muciniphila*, *Bacteroides thetaiotaomicron, Bacteroides caccae, Bacteroides ovatus, Bacteroides uniformis, Barnesiella intestinihominis, Clostridium symbiosum, Collinsella aerofaciens, Desulfovibrio piger, Escherichia coli, Marvinbryantia formatexigens, Agathobacter rectalis, Facecalibacterium prausnitzii* and *Roseburia intestinalis*. AEX= anion exchange chromatography, LC= liquid chromatography. ‘+’ = positive samples containing phage, ‘-’ = negative control samples not containing phage.

While we detected most of the bacterial community members (11/14) in the pre-DNase mastermix sample, the spike-in phage DNA made up a far larger portion (Fig. S1) likely due to the removal of bacterial cells in the initial centrifugation and filtration steps^18,34,35^. We detected the missing members in extremely low abundances at later stages of the purification (Fig. 2B, Suppl. Datasets 1-2). Additionally, while all five spike-in phage were detected (Fig. S1), M13 was detected at a relatively low abundance despite adding an equal number of plaque forming units (PFUs) of each phage. This is in part due to the small genome size of M13, which leads to a smaller number of sequencing reads and in part to its ssDNA genome which does not sequence well with Illumina platforms unless special library preparation kits are used. After treatment with DNase, most of the bacterial DNA was reduced, however it was not completely removed as is expected based on prior literature^18,36^ (Fig. S1). Interestingly, the DNA of certain bacterial taxa, *Desulfovibrio piger*, *Clostridium symbiosum* and *Marvinbryantia formatexigens,* seemed to be less efficiently removed than DNA from other taxa suggesting that it may have been protected in VLPs. Finally, while most of the phage were not affected by the DNase digestion, M13 abundance was drastically reduced following digestion suggesting that DNase, the incubation, or the combination of the two may bias against filamentous phage (Fig. 3B). Had we not added M13 in the quantity we did (1e9 PFUs per replicate), it may have been completely lost during digestion.

**Figure 2.**
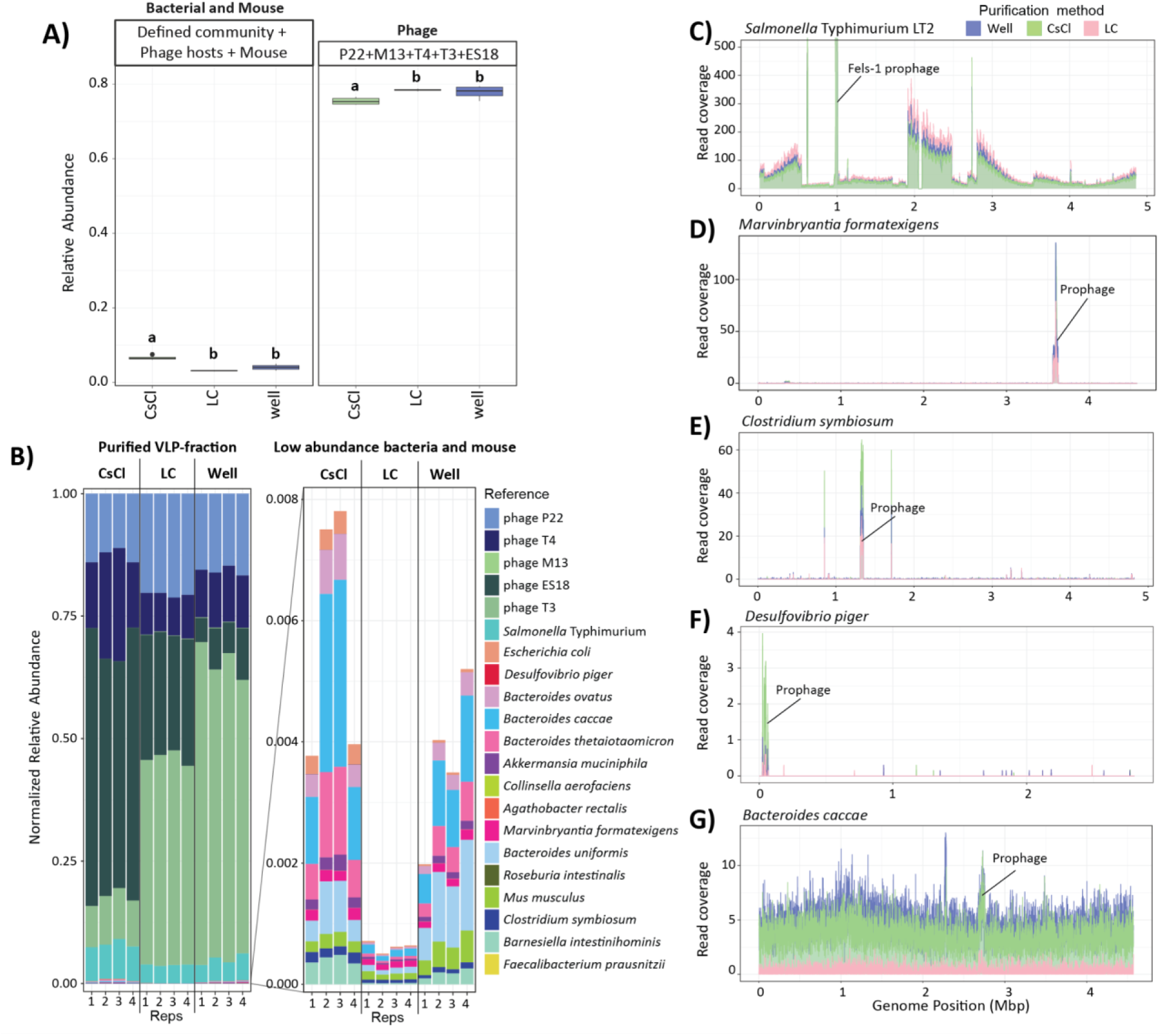
Comparison of the LC, 24-Well and CsCl purification methods. Relative abundances were determined by mapping reads to the reference genomes for the phage, bacteria and mouse and dividing the number of assigned reads by the total number of trimmed and decontaminated reads for the respective sample. A) Relative abundance of bacterial (All defined community members and hosts used for phage propagation) and phage (P22, ES18, T3, T4 and M13) DNA in LC, Well and CsCl purified samples. Statistical significance was determined with a beta generalized linear mixed-effects model (p<0.05). If different letters (a,b) appear above two box plots this indicates that they are significantly different. B) Taxonomic composition of LC, CsCl and 24-Well purified VLP-fractions. Relative abundances were normalized to 1 by dividing the relative abundance values by the sum of relative abundances in each sample. Visualization of read coverage along the genomes of C) *Salmonella enterica* serovar Typhimurium, D) *Marvinbryantia formatexigens*, E) *Clostridium symbiosum*, F) *Desulfovibrio piger* and G) *Bacteroides caccae*. All genome coverage plots use the coverage data from replicate 4.

**Figure 3.**
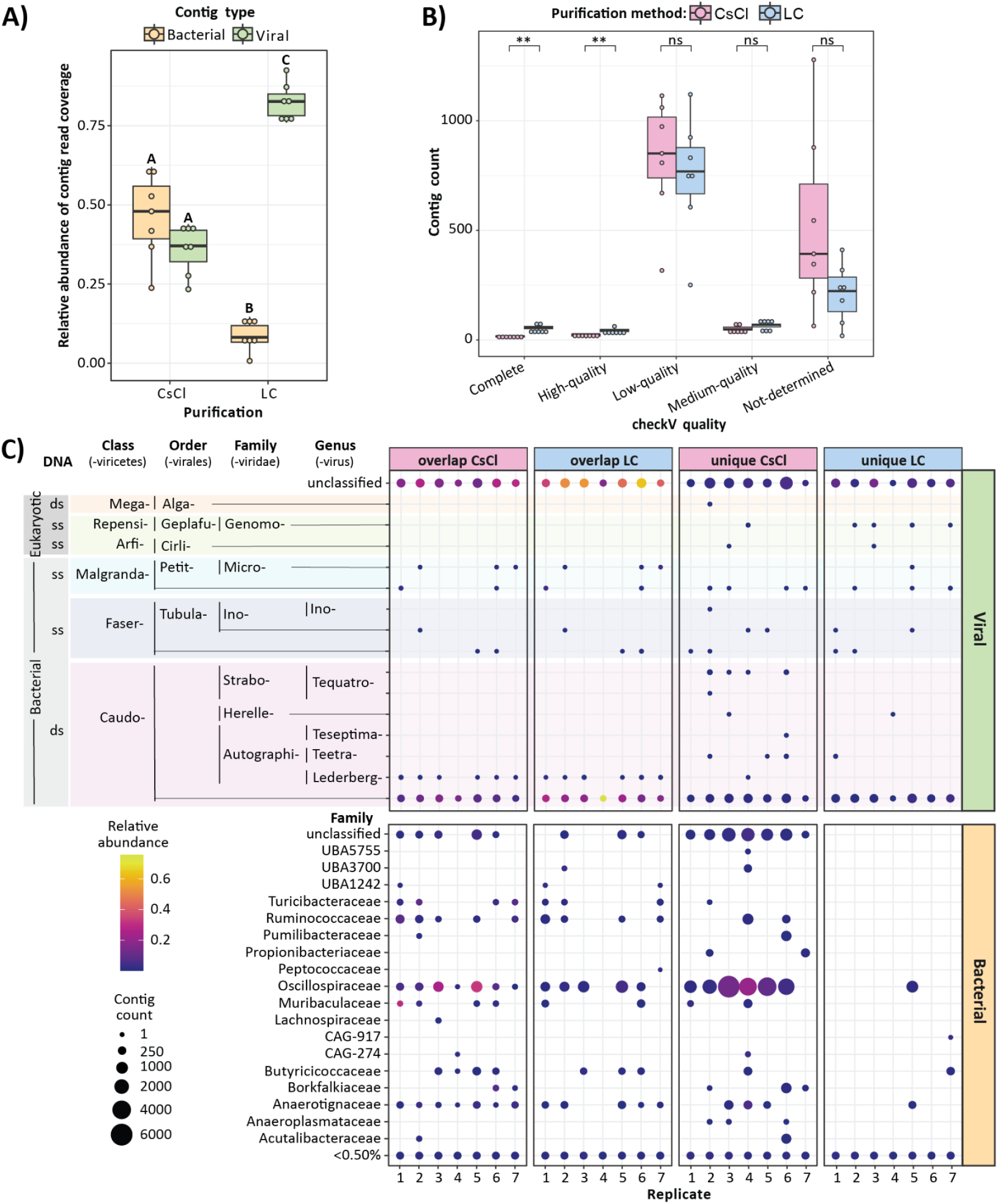
Metagenomic characterization of VLPs purified with AEX-LC and CsCl density gradients from fecal pellets of mice with a conventional microbiota. A) Relative abundance of reads mapping to bacterial and viral contigs in the CsCl density-gradient purified samples (CsCl) (36% avg. viral, 46% avg. bacterial) and the anion exchange liquid chromatography (LC) (82% avg. viral, 9% avg. bacterial) purified samples. Contigs were assembled from shotgun metagenomic sequencing reads of VLP-fractions and classified into bacterial and viral contigs using four different viral classification tools. Contigs not classified as viral by at least two tools or contigs containing a prophage were considered bacterial. Relative abundance was determined by mapping reads to contigs and dividing by the number of trimmed and decontaminated reads for the respective sample. Statistical significance of differences between bacterial and viral contig abundances both between and within conditions is indicated with letters (A,B,C) above box plots (beta generalized linear mixed model (GLMM), p<0.05). If letters are different then a comparison is significantly different. B) The checkV quality of viral contigs in the CsCl and LC purified samples. There were significantly more complete and high quality viral genomes assembled in the LC purified samples compared to the CsCl samples (t-tests, p<0.05). C) Comparison of bacterial and viral contigs that are unique and shared amongst LC and CsCl purification methods. Size of circle represents the number of contigs associated with each bacterial or viral taxonomic group and color of circle represents summed relative read abundance of the associated contigs. Relative abundance was determined by summing the reads that mapped to contigs associated with a specific taxonomic group then dividing by the number of trimmed and decontaminated reads for the respective sample.

#### VLPs purified using AEX are composed of less bacterial DNA and more viral DNA compared to VLPs purified with CsCl

By collecting fractions throughout the AEX-based purifications, we were able to determine if VLPs and free DNA flowed through during loading (Load), bound and eluted during the elution step (Elute), or bound so strongly that they only eluted during the clean-in-place step (CIP). For both LC and Well purifications, the phage had the highest relative abundances in the Elute fractions with one exception. Phage ES18 had a higher relative abundance in the Well CIP fractions suggesting that it may not have eluted well compared to the other phage (Fig. S2). Overall, the proportion of bacterial DNA in the Well and LC Elute fractions was significantly lower than the proportion of bacterial DNA in the CsCl-purified VLPs (beta generalized linear mixed model (GLMM), p<0.05) (Fig. 2A). Additionally, while all the spike-in phage were captured by each of the three purification methods, the proportions were different, highlighting potential biases between methods (Fig. 2B). CsCl purification captured relatively more ES18 phage than LC or Well-purified VLPs, however phage M13 was almost completely lost in CsCl purification potentially due to the known bias of the CsCl method against more fragile phage morphologies (Fig. 2B, Fig. S3)^23^.

For both AEX-based methods, most of the bacterial DNA was in the Load or CIP fractions, while the phage were primarily in the Elute fraction (Fig. S2). Interestingly, the DNA of three bacteria - *S. enterica* sv. Typhimurium LT2, *C*. *symbiosum* and *M. formatexigens* had relatively higher abundances in the LC Elute fraction compared to the other bacteria (Fig. S2). The DNA from these bacteria was also less efficiently digested by DNase relative to the other taxa (Fig. S1). We investigated the genome coverage of these taxa and found that the *S. Typhimurium* DNA enrichment in the VLP fraction was partially a result of DNA packaging into phage P22 during generalized transduction which generates a specific read coverage pattern^11,37^ and partially a result of strong Fels-1 induction, a resident prophage in the LT2 genome (Fig. 2C). The *C. symbiosum* and *M. formatexigens* DNA in the Elute fractions were also results of resident prophage induction (Fig. 2D, Fig. 2E). We also found these prophages in the CsCl purified VLPs (Fig. 2D, Fig. 2E). These prophages are present at extremely low abundances demonstrating each purification method’s ability to capture rare, low abundance VLPs (Fig. 2B, Fig. S3, Suppl. Datasets 1-2). For example, phage P22 was present in the purified VLPs at a relative abundance of ∼10-16%, while the prophage in *M. formatexigens* was present at ∼0.009%-0.018% relative abundance. We inspected the genome coverage of the other bacterial taxa detected in the purified VLP fractions and found that *D. piger* also had an induced prophage which is likely why its abundance did not decrease during DNase digestion (Fig. 2F, Fig. S1). Some of the bacterial taxa had relatively full, albeit low, genome coverage with occasional spikes in coverage associated with induced prophage while others had sparse to no coverage (Fig. S4). *B. caccae*, for example, had an induced prophage however the signal was partially masked by the background coverage across the genome (Fig. 2G). The full genome coverage of certain bacterial taxa may indicate contamination or the presence of transducing VLPs that package the whole host bacterial genome such as GTAs or MVs^38–41^.

### VLPs purified from fecal pellets of healthy mice with a conventional microbiota

#### LC-purified VLPs are composed of significantly more viral DNA compared to CsCl purified VLPs

To demonstrate the application of AEX-LC VLP ultra-purification to a relevant complex microbiome, we purified VLPs from conventional murine fecal homogenates (n=7) with AEX LC and CsCl. The sequencing reads from the purified VLPs were assembled and resulting contigs classified as viral with geNomad^42^, VirSorter2^43^, VIBRANT^44^ and DeepVirFinder^45^ (Fig. S5, Fig. S6). Contigs were considered viral if they were classified as viral by two or more tools. There were a total of 41,228 contigs >2 kbp assembled in the LC VLP-fractions, of which 16,132 were classified as viral by at least one tool and 8,095 were classified as viral by two or more tools. In the CsCl VLP-fractions, there were a total of 68,207 contigs >2 kbp assembled of which 29,658 were classified as viral by at least one tool and 10,627 were classified as viral by two or more tools. All viral contigs were run through CheckV^46^ to determine both viral genome quality and prophage status. If the viral contig was classified as a prophage by CheckV (e.g. if the viral portion was flanked by bacterial genes on both the 3’ and 5’ ends), it was re-classified as bacterial (Fig. S7). We determined viral contig taxonomy using the IMG/VR database^47^ and bacterial contig taxonomy with CAT^48^. The majority of viral (73%) and bacterial (97%) contigs received some level of taxonomic classification. We calculated the relative abundance of all bacterial and viral contigs in both the CsCl and LC purified VLP metagenomes based on mapping of sequencing reads and determined that there was a significantly higher (GLMM, p<0.05) proportion of viral DNA in the LC purified VLP-fractions (85%) compared to the CsCl purified VLP-fractions (41%) (Fig. 3A). While the differences were small, there were significantly more complete and high quality viral genomes assembled in the LC purified samples compared to CsCl (t-test, p<0.05) (Fig 3B).

#### LC-purified VLPs capture a similar viral profile as CsCl-purified VLPs

We compared the similarities and differences of contigs assembled from LC and CsCl purified VLPs using Redundans^49^ to find overlapping contigs between assemblies (Fig. 3C). The approach we used allows for large contigs to have multiple smaller contigs overlapping and therefore there are occasionally differences in contig counts between overlapping contigs (Fig. 3C). Additionally, a small percentage (0.018%) of overlapping contigs did not have the same classification as either viral or bacterial. For these, we used the classification and associated taxonomy of the larger contig. Out of 18,349 viral contigs assembled in total, there were 6,436 and 4,481 viral contigs unique to CsCl and LC purified VLPs, respectively, while 7,432 viral contigs were shared between purification methods. However, the unique viral contigs summed to only small relative read abundances (avg. 4% in CsCl VLPs, avg. 8% in LC VLPs) and therefore lead to small changes in overall VLP-fraction composition. The overlapping viral contigs had far larger relative abundances (32% avg. in CsCl and 73% avg. in LC). The most abundant viral taxonomic groups overall in each purification method were unclassified Caudoviricetes (i.e. tailed phage) (avg. 12% in CsCl and 30% avg. in LC) and completely unclassified viruses (avg. 24% in CsCl and avg. 50% in LC). These viral groups were largely shared between the LC and CsCl purified samples (Fig. 3C).

The Genomoviridae (eukaryotic viruses) and the Tequatroviruses contigs were consistently unique to the LC and CsCl VLP-fractions, respectively, however they were each represented by very few contigs (1-2 in LC reps, 1-28 in CsCl reps) (Fig. 3C). There were also a large number of unclassified viral contigs that are unique to each purification method (Fig. 3C). The biases between purification methods may be due to several factors. For the AEX LC method these include the fact that positively charged VLPs will not bind to an AEX column and would therefore be lost during LC purification, however previous literature has shown that most phage have negative surface charge^50,51^. Additionally, VLP binding may be impacted if the sample is not the same pH and conductivity as the loading buffer. We diluted our samples with the LC loading buffer to achieve similar pH and conductivity values, however, dialysis or filter-based methods can achieve closer matches between values albeit at the expense of increased processing times and potential sample loss. Biases in the CsCl density gradient centrifugation method may be due the fact that it is a physically harsh method that can bias against structurally weak or fragile VLPs^23^. Additionally, VLPs that have a buoyant density <1.35 g/mL will be lost as they will not reach the 1.35/1.5 g/ml interface. To alleviate this, a lower density layer would need to be used, which would increase risk of contamination or multiple density layers would need to be extracted for sequencing. In summary, while there were differences in VLPs purified by each purification method, likely due to differences in VLP properties used for purification (charge vs buoyant density), the overall profile of abundant viruses was largely similar between methods.

#### Specific bacterial families are represented in the bacterial DNA portion of murine fecal VLP-fractions

The VLP-fractions of both LC and CsCl purification methods showed a consistent presence of DNA from several bacterial families and genera (Fig. 3C, Fig. S8). One of the major differences between the LC and the CsCl method was that CsCl yielded a higher total number of bacterial contigs (69,403 in CsCl vs 40,978 in LC) and had more unique bacterial contigs (47,896) compared to the LC (19,741) assemblies. Additionally, contigs from specific bacterial genera were much more abundant in the CsCl method than in the LC method (Fig. S8) corresponding with the much higher overall abundance of bacterial DNA in the CsCl VLP-fractions (Fig. 3A). The Oscillospiraceae, Anaerotignaceae, and Ruminococcaceae were among the families that were most consistently present and had the most contigs. The Oscillospiraceae stood out in particular due to the much higher number and relative abundance of contigs in the CsCl purified VLP-fractions compared to the LC purified VLP-fractions. This data is in line with previous observations of high amounts of Oscillospiraceae derived DNA in CsCl purified VLPs^52,53^. Currently we do not know what causes this “enrichment” of Oscillospiraceae in CsCl purified VLPs. One possible explanation for the bacterial DNA in the purified VLP-fractions is that the DNA originates from small bacterial cells that pass the 0.45 µm filter and were co-purified with the VLPs. An alternative explanation is that the abundant DNA in CsCl purified VLPs is derived from abundant transducing VLPs or MVs that carry bacterial host DNA. For example, the Oscillospiraceae and Ruminococcaceae have been implicated in GTA-mediated transduction mechanisms which are associated with the packaging and transfer of large regions of the bacterial genome^45^, however, it is unclear if GTAs alone could explain the large quantities of VLP carried DNA. An additional mechanism for VLP carriage of large bacterial genome regions are MVs ^38,39,55^ which are often co-purified with VLPs in density gradients due to their similar buoyant densities^56,57^.

## Conclusions

The challenges associated with CsCl purifications are a barrier for widespread and large-scale viromics research. Here, we show that AEX chromatography is a reliable alternative to the gold standard CsCl method for VLP purification prior to DNA sequencing. AEX chromatography significantly reduces sample processing time, allows for automation when FPLC is used and parallelized processing of large sample numbers when multi-well plates are used, and improves viral purity compared to CsCl purifications. The proportion of bacterial DNA in the VLPs purified by both the LC and Well chromatography methods was consistently lower than in the CsCl-purified VLPs. While chromatography-purified VLPs were not devoid of bacterial DNA, we found that in gnotobiotic mouse samples with phage spike-ins, much of the bacterial DNA was associated with carriage of bacterial DNA in VLPs (Fig. 2). Similarly, we found evidence of VLP-packaged bacterial DNA in conventional murine fecal microbiomes, where the abundance of bacterial DNA was much higher in the CsCl-purified VLPs as compared to the LC-purified VLPs. Ultimately, however, future work using approaches such as electron microscopy and/or proteomics will be needed to definitively determine if this bacterial DNA is packaged into VLPs or free.

While the LC method has the advantage of automation using autosamplers and automated fraction collectors, the Well method has a throughput advantage. A standard table-top centrifuge with a 4-bucket swing-out rotor can be used to process 4 x 24-well plates (96 samples) in parallel thereby significantly increasing throughput and making large-scale viromics experiments feasible. AEX multi-well plates also exist in 96-well format, which allows for increased throughput, however the volume that can be loaded in each well is much smaller, making use of 96-well plates only applicable if sample concentrations are very high and thus required loading volumes low. Interestingly, we did notice differences between the LC and Well-purified VLP-fractions despite the use of the same AEX column material and purification method. For example, the ES18 phage did not elute off the Well columns as efficiently as it eluted off the LC column and conversely, the M13 phage was present at higher relative abundances in the Well Elute fractions compared to the LC. This variability may be due to differences in processing caused by the fact that the Well plate was loaded, washed, eluted, and CIPed with multiple separate loads that were completely spun through the column material, while in the LC method a continuous flow of buffer occurs and the column never falls dry during sample purification.

Future work can use diverse monolithic column types, buffer systems and elution gradients to optimize VLP purification methods for studying viromes from specific environments or to better enrich specific viruses or VLP types. For example, monolithic hydrophobic interaction (HIC) and multi-modal columns (combining AEX and HIC) have both been successfully used for single VLP purifications^28,58^. Additionally, the buffer system used including the pH and elution salt can have a major effect on the VLPs and/or debris retained on an ion exchange column and therefore testing different buffer systems may result in the capture of different VLPs. Furthermore, while we used an isocratic elution for the murine fecal VLP LC purifications, an elution gradient may be able to separate VLPs with specific characteristics from each other thus allowing for greater control over the associated viromics study^59^. Finally, taking samples for sequencing pre- and post-DNase digestion or throughout the chromatography purification and comparing relative abundances of bacterial species can be used to identify bacteria that are producing VLPs, even at low abundances.

## Methods

### Generation of fecal homogenates from gnotobiotic mice with phage spike-ins

#### Growth of bacteriophages for spike-ins

P22: We diluted an overnight culture of *Salmonella enterica* serovar Typhimurium LT2 (LT2) grown in LB media (1% (g/L) Tryptone, 0.05% Yeast, 1% NaCl) to 0.4 OD 600 nm (OD) and infected with P22 at a multiplicity of infection (MOI) of 0.01 and incubated at 37°C while shaking for 5-6 hours.

M13: We diluted an overnight culture of *Escherichia coli* 2191 (2191) grown in LB media to 0.4 OD and infected with M13 at a MOI of 0.4 and incubated at 37°C while shaking for 24 hours.

ES18: We diluted an overnight culture of LT2 grown in LB media to 0.4 OD and infected with ES18 at a MOI of 0.01 and incubated at 37°C while shaking for 5 hours.

T4: We diluted an overnight culture of *Escherichia coli* C (ECC) grown in LB media to 0.4 OD and infected with T4 at a MOI of 0.4 and incubated at 37°C while shaking for 5 hours.

T3: We diluted an overnight culture of ECC grown in LB media to 0.4 OD and infected with T3 at a MOI of 0.001 and incubated at 37°C while shaking for 3 hours.

#### Clarification of bacteriophages to be used as spike-ins

We centrifuged each of the phage lysates at 12,000 x g for 15 minutes to pellet cell debris. The supernatants were removed and filtered using 0.45 µm bottle top filters (Genesee Scientific, catalog no. 25-234). The filtered lysates were titered using the drop-plate method.

#### Generation of gnotobiotic fecal pellets with phage spike-ins

Germ-free C57BL/6J mice were gavaged with a defined microbiota consisting of 14 members - *Akkermansia muciniphila* 22959*, Bacteroides ovatus* 1896*, Bacteroides thetaiotaomicron* 2079*, Bacteroides caccae* 19024*, Bacteroides uniformis* 8492*, Barnesiella intestinihominis* YIT11860*, Clostridium symbiosum* 934*, Collinsella aerofaciens* 3979*, Desulfovibrio piger* 29098*, Escherichia coli* HS*, Marvinbryantia formatexigens* 14469*, Agathobacter rectalis* 33656 *(Eubacterium rectale), Facecalibacterium prausnitzii* 17677 and *Roseburia intestinalis* 14610^33,60^. Mice were held in gnotobiotic isolators for 9 months after gavage after which fecal pellets were sampled and frozen at −80°C.

A fecal homogenate was created by manually homogenizing 19 gnotobiotic murine fecal pellets in 19 mL of LC/24-well loading buffer (50 mM HEPES, pH 7.5) using sterile inoculating loops. The homogenate was centrifuged at 2,000 x g for 10 minutes to pellet larger debris. The supernatant was removed and centrifuged again at 5,000 x g for 10 minutes. The supernatant was filtered using a 0.45 µm syringe filter (VWR, catalog no. 76479-020) and split into a positive and negative control aliquot. We added equal amounts of PFUs (1.6×10^10^ PFUs per phage) of each clarified spike-in phage to the positive control aliquot to generate a master mix. We treated the positive and negative control aliquots with 50 U/mL of DNase 1 (Thermo Scientific, catalog no. PI90083) and incubated at 37°C for 2 hours. We split the positive and negative controls again into replicates for purification with each of the purification methods (12x positive aliquots in total, 4x negative aliquots in total). We estimated that each replicate would contain roughly 10^9^ PFUs of each phage. With this PFU number per sample we accounted for controls and sequencing aliquots to be taken throughout the experiment. See figure 1 for experimental overview.

### Fecal homogenate from mice with a conventional microbiota

#### Sampling of mouse fecal material from mice with a conventional microbiota

Seven C57BL/6J mice (male, 4-5 weeks old) purchased from Jackson Laboratories were used to obtain fecal pellets for comparing the CsCl and the LC methods using complex microbiome samples. Mice were housed with autoclaved bedding and water, and irradiated standard chow. All mice were subjected to a 12-hour light and 12-hour dark cycle. Experiments were conducted in the Laboratory Animal Facilities located on the NC State CVM campus. The animal facilities are equipped with a full-time animal care staff coordinated by the Laboratory Animal Resources (LAR) division at NC State. The NC State CVM is accredited by the Association for the Assessment and Accreditation of Laboratory Animal Care International (AAALAC). Trained animal handlers in the facility fed and assessed the status of animals several times per day. Those assessed as moribund were humanely euthanized by CO_2_ asphyxiation. This protocol is approved by NC State’s Institutional Animal Care and Use Committee (IACUC).

Two fecal pellets were collected per mouse. The pellets were manually homogenized in 1.2 mL of SM buffer (100 mM NaCl, 8 mM MgSO^4-^*7H_2_O, 50 mM TrisHCl) using sterile inoculating loops. After homogenization, the samples were brought up to 2 mL total volume using SM buffer. An aliquot of each replicate’s homogenate was removed and kept at −80°C for whole-genome sequencing.

Fecal homogenates were centrifuged at 2,500 x g for 5 minutes and the supernatants were transferred to clean tubes. Samples were centrifuged again at 5,000 x g for 5 mins to pellet residual debris and supernatants were removed and filtered using 0.2 µm filters (VWR, catalog no. 76479-020) into clean tubes. 50 U/mL DNase 1 (Thermo Scientific, catalog no. PI90083) was added to each tube and samples were incubated at 37°C for 2 hours, after which samples were split 1:1 and VLPs ultrapurified using either AEX FPLC or CsCl.

### Purification by AEX using an FPLC or 24-well plates

We used an ÄKTA Pure chromatography system and a monolithic 1 mL CIMmultus QA-AEX column (Sartorius #: 311.5113-2) with 2 µm pores for our FPLC-based purifications. We used a 50 mM HEPES, pH 7.0 buffer system with 1 M NaCl added for elution. Each replicate sample was diluted 1:7 with loading buffer prior to sample loading. The instrument and column were equilibrated with 10 columns volumes (CVs, 1 CV = 1 ml)) loading buffer, 10 CVs elution buffer, and another 10 CVs of loading buffer before each replicate was loaded. After loading the sample, the column was washed with 25-30 CVs of loading buffer. The sample was eluted isocratically with 15 CVs of 1 M NaCl followed by clean in place (CIP) with 15 CVs of 1 M NaOH, 2 M NaCl solution. For the spike-in samples the last 2mL of loading and the first 2 mL of elution and CIP were collected as fractions into separate collection tubes using an autosampler (Fig. S9). The fractions were titered using the drop-plate method on bacterial hosts LT2, 2191, and ECC. Fractions were washed 3x with SM buffer and concentrated to ∼500uL using 10 kDa MWCO centrifugal filters (Amicon, Millipore Sigma, catalog no: UFC8010). For the homogenate from conventional mouse fecal pellets the first 2 ml of the elution were collected in an autosampler tube (Fig. S10).

Purification using 24-well plates replicated the FPLC method. The 24-well plates contained 1 mL CIMmultus QA-AEX monolithic chromatographic media (Sartorius #: BIA-124.5113-2) with 2 µm pores (1 CV = 1 ml). We centrifuged the plate at 750 x g for 2 minutes for each step (equilibration, loading, elution and CIP) in the purification. Since the plate allows for addition of 6 ml working volume, some of the steps were carried out by multiple sequential centrifugations to replicate the CVs used for each step of the FPLC method. 2 mL fractions were collected during sample loading, elution and CIP corresponding to the same CVs as in the FPLC method. The fractions were titered on bacterial hosts LT2, 2191, and ECC and washed 3x with SM buffer and concentrated to ∼500uL using 10 kDa MWCO centrifugal filters.

### CsCl purification

We prepared density gradients using three cesium chloride (CsCl) density layers- 1.7 g/mL, 1.5 g/mL and 1.35 g/mL- in Beckman ultra-clear tubes (14×89 mm, catalog no. 344059). The interface between the 1.35/1.5 g/mL layers was marked on the tube for each replicate. For the spike in samples 1 mL of sample was loaded onto each density gradient. For the conventional mouse fecal homogenate 750 µL were loaded. The samples were centrifuged at 70,000 x g for 24 hours. After centrifugation, the 1.5/1.35 g/mL interface was extracted by piercing the tube at the interface using a syringe needle. The 1.5/1.35 g/mL interface is enriched with VLPs^61^. The resulting sample was washed 3x with SM buffer and concentrated to ∼500uL using 10 kDa MWCO centrifugal filters (Amicon, Millipore Sigma, catalog no: UFC8010).

### DNA extraction and sequencing

Samples were treated 1:1 with lysis mix (2% SDS, 90 µg/mL Proteinase K) and incubated at 55°C for 1 hour. Samples were cooled to room temperature and transferred to a pre-spun 5-prime light phase lock tube (QuantaBio, catalog no: 2302820) for phenol:chloroform DNA extraction. 0.5 mL of phenol:chloroform:isoamyl (25:24:1) was added to each tube and inverted to mix. Tubes were centrifuged at 12,000 x g for 5 minutes. The aqueous phase was poured off into a new phase-lock tube. 0.5 mL of chloroform:isoamyl (24:1) was added to each tube and inverted to mix. Tubes were centrifuged at 12,000 x g for 5 minutes. The aqueous phase was poured off into clean microcentrifuge tubes. The extracted DNA was cleaned and concentrated using Qiagen MinElute columns (Qiagen, catalog no: 28004). Samples were diluted 5:1 in the provided Buffer PB and loaded onto the spin-column for centrifugation. The flow through was discarded and 750 µL of the provided buffer PE was added to the spin-column and centrifuged. The flow through was discarded and the spin-columns were centrifuged. The spin-columns were placed into clean tubes and 20 µL of the provided buffer EB was added to the membranes and centrifuged to elute the DNA. All centrifugation was performed at 17,900 x g for 1 minute. DNA concentrations of all samples were determined using a Qubit HS DNA (Thermo Fisher, catalog no: Q32851) assay prior to sequencing. Sequencing libraries were prepared with Twist Biosciences enzymatic fragmentation kit and UDI/UMI primers. Samples were sequenced on the Illumina NovaSeq X platform at 2×150 bp. We requested 50 million reads per sample, however due to large differences in DNA input between samples (e.g. loading fractions in FPLC were very low), we received between 8.8 M and 150 M paired-end reads for the spike-in samples. Importantly, for the spike-in samples the key samples for each of the methods (elution fractions for FPLC and 24-well plate) all had around 50 million paired-end reads making them comparable across methods. For the fecal homogenates from mice with a conventional microbiota, sequencing libraries were prepared using the xGen DNA library prep kit (IDT) and xGen normalase and UDI primers. Samples were sequenced on the Illumina NovaSeq X platform at 2×150 bp. We requested 150 million reads per sample and we received between 26 M and 486 M paired-end reads.

### Data analysis

#### Fecal spike-in sample

Raw sequencing reads were trimmed with TrimGalore to remove Illumina adapter sequences and any overrepresented sequences (e.g. PolyGs and Ns, determined via FastQC) (Fig. S11) and mapped to reference genomes which included the bacteria gavaged into the mice, the bacterial hosts of the phage used as spike-ins, the phage, and the mouse genome (Table S2) using BBMap^62^ (ambiguous=random, minid=0.97, and refstats output).

Nearly 80% of reads mapped to the reference genomes in the key samples for each of the methods (elution fractions for FPLC and 24-well plate) (Fig. S12). Relative abundance was determined by dividing the number of reads assigned to a specific genome by the total number of trimmed reads for the respective sample. While there were three strains of *E. coli* present in our samples (*E. coli* C and E. coli 2191 used for phage propagation and *E. coli* HS in the 14-member bacterial community), we only used the *E. coli C* genome for read mapping to avoid redundancy issues that might affect abundance calculations. Pileup files were generated with BBMap’s^62^ pileup.sh (binsize=1000) for visualizations of bacterial genome coverage. We used geNomad’s^42^ end-to-end pipeline with score calibration to predict prophage locations in bacterial genomes.

#### Fecal homogenates from mice with a conventional microbiota

Raw sequencing reads were trimmed using TrimGalore. Reads originating from the mouse or phiX phage were removed using BBsplit (ambig=split) with the mouse and phiX genomes (GCF_000001635.27 and NC_001422.1, respectively) as references (Fig. S13). Reads originating from each sample were individually assembled using MEGAHIT V1.2.9^63^ with default parameters. We used geNomad^42^ (end-to-end, -enable-score-calibration), VirSorter2^43^, VIBRANT^44^ (virome, length=2000) and DeepVirFinder^45^ (length=2000) to identify viral contigs in each assembly. We used the ‘Viral Sequence Identification with VirSorter2 V.3^64^’ pipeline for VirSorter2. This pipeline included running VirSorter2 (include-groups=dsDNAphage, ssDNA, min-length=2000, min-score=0.5), feeding the results into checkV to identify viral and prophage containing contigs, and feeding those contigs into VirSorter2 again (viral-gene-enrich-off, include-groups= dsDNAphage, ssDNA, min-length=2000, min-score=0.5)). A contig was considered viral if at least two tools agreed on the viral classification. We ran checkV^46^ on the final viral contig set to identify prophages which were re-classified as ‘bacterial’ DNA. We used CAT^48^ to determine the taxonomy of the bacterial contigs. We determined the taxonomy of viral contigs by searching against the IMG/VR database^47^ with BLASTn^65^ (minimum sequence id= 50%, minimum query overlap=50%). Relative abundance was determined by mapping reads to the contigs with BBMap (ambiguous=random, minid=0.97) (∼80% of reads mapped to the assemblies (Fig. S14)), dividing the number of assigned reads by the total number of trimmed and decontaminated reads for the respective sample. We used Redundans^49^ (minimum sequence id=95%, minimum query overlap=95%) to determine overlap of viral contigs between assemblies.

## Supplemental files

**Supplemental figures and tables:** SupplementaryMaterials

**Supplemental datasets:** DatasetS1, DatasetS2

## Acknowledgements

The authors would like to thank Casey Theriot, Karen Flores and Ayesha Awan for collecting fecal pellets for this work and John Van Zaten for providing input for LC purifications. The authors would also like to thank all members of the Kleiner Lab for their consistent feedback and support throughout this work.

## Funding

This research was supported by a seed grant from the North Carolina State University Data Science Academy and by the National Institutes of Health under Award Numbers R35GM138362 and R01Al171046. The content is solely the responsibility of the authors and does not necessarily represent the official views of the National Institutes of Health.

## Contributions

**Jessie Maier:** Conceptualization, Data Curation, Formal Analysis, Investigation, Methodology, Visualization, Writing- original draft, Writing- editing and review

**Namita Deshmukh:** Investigation, Methodology

**Manuel Kleiner:** Conceptualization, Funding Acquisition, Resources, Writing- editing and review

## Data availability

**Gnotobiotic fecal samples with phage spike-ins:**

The raw sequence datasets are available in the Sequence Read Archive (SRA) repository under BioProject PRJNA1482603.

**Conventional mouse fecal VLPs:**

The raw sequence datasets are available in the Sequence Read Archive (SRA) repository under BioProject PRJNA1404639.

## Notes

### Competing Interest Statement

The authors have declared no competing interest.

