## SupplementalMaterials for "High throughput chromatographic ultra-purification of virus-like particles for downstream viromics"

**Affiliations:**
[1]Department of Plant and Microbial Biology, North Carolina State University, Raleigh, NC

Supplementary Figures:

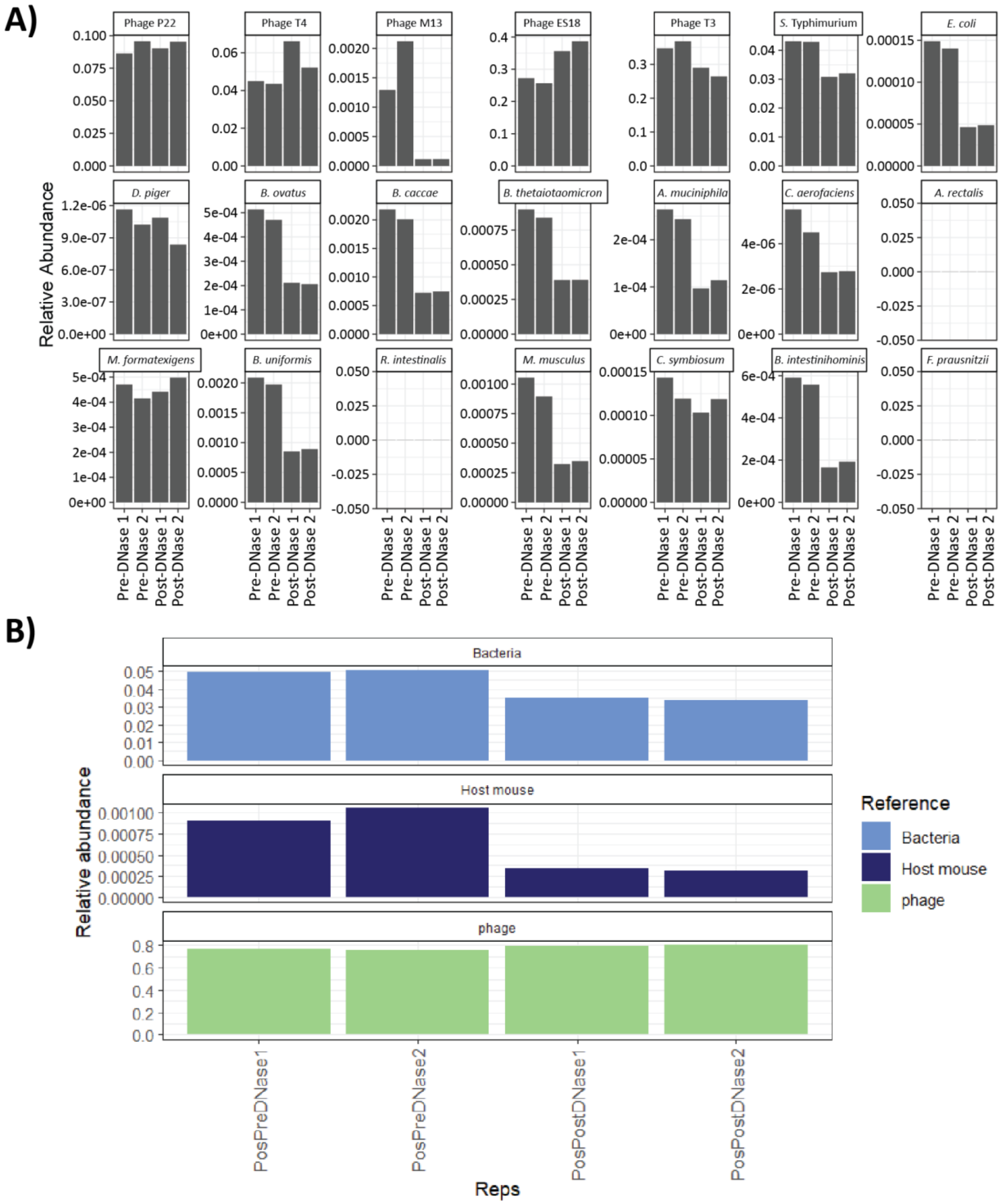


**Supplementary figure 1 - Relative abundances of phage and bacterial taxa in the gnotobiotic mouse fecal samples before and after DNase treatment.** DNase treatment was performed with 50 U/mL DNase for 2 hours at 37 °C. Two technical replicates of each condition were taken for DNA extraction and sequencing. A) Relative abundance of each individual phage, bacterium and host mouse genome in the sample pre- and post- DNase treatment. B) Relative abundance of summed phage, bacterial and house mouse DNA pre- and post- DNase treatment. Full names of bacterial taxa are as follows- *Akkermanisa muciniphila*, *Bacteroides thetaiotaomicron*, *Bacteroides caccae*, *Bacteroides ovatus*, *Bacteroides uniformis*, *Barnesiella intestinihominis*, *Clostridium symbiosum*, *Collinsella aerofaciens*, *Desulfovibrio piger*, *Escherichia coli*, *Marvinbryantia formatexigens*, *Agathobacter rectalis*, *Facecalibacterium prausnitzii* and *Roseburia intestinalis*.

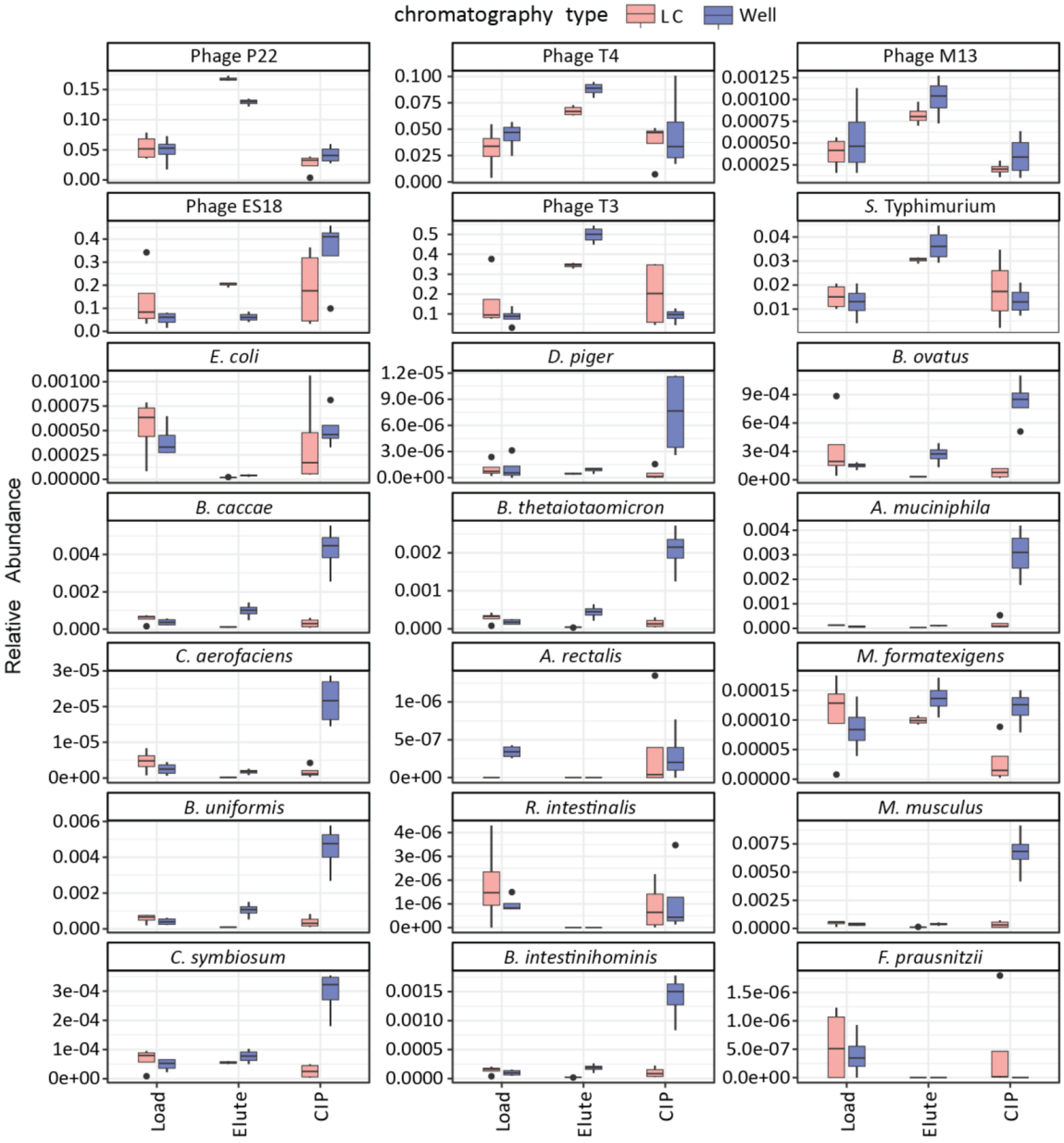

**Supplementary figure 2 - Relative abundance of phage and bacterial taxa in the gnotobiotic mouse fecal samples during sample load (Load), elution (Elute) and clean-in-place (CIP) for both anion exchange (AEX) liquid chromatography (LC) and 24-well assay plates with AEX column material (Well).** Full names of bacterial taxa are as follows- *Akkermanisa muciniphila*, *Bacteroides thetaiotaomicron*, *Bacteroides caccae*, *Bacteroides ovatus*, *Bacteroides uniformis*, *Barnesiella intestinihominis*, *Clostridium symbiosum*, *Collinsella aerofaciens*, *Desulfovibrio piger*, *Escherichia coli*, *Marvinbryantia formatexigens*, *Agathobacter rectalis*, *Facecalibacterium prausnitzii* and *Roseburia intestinalis*.

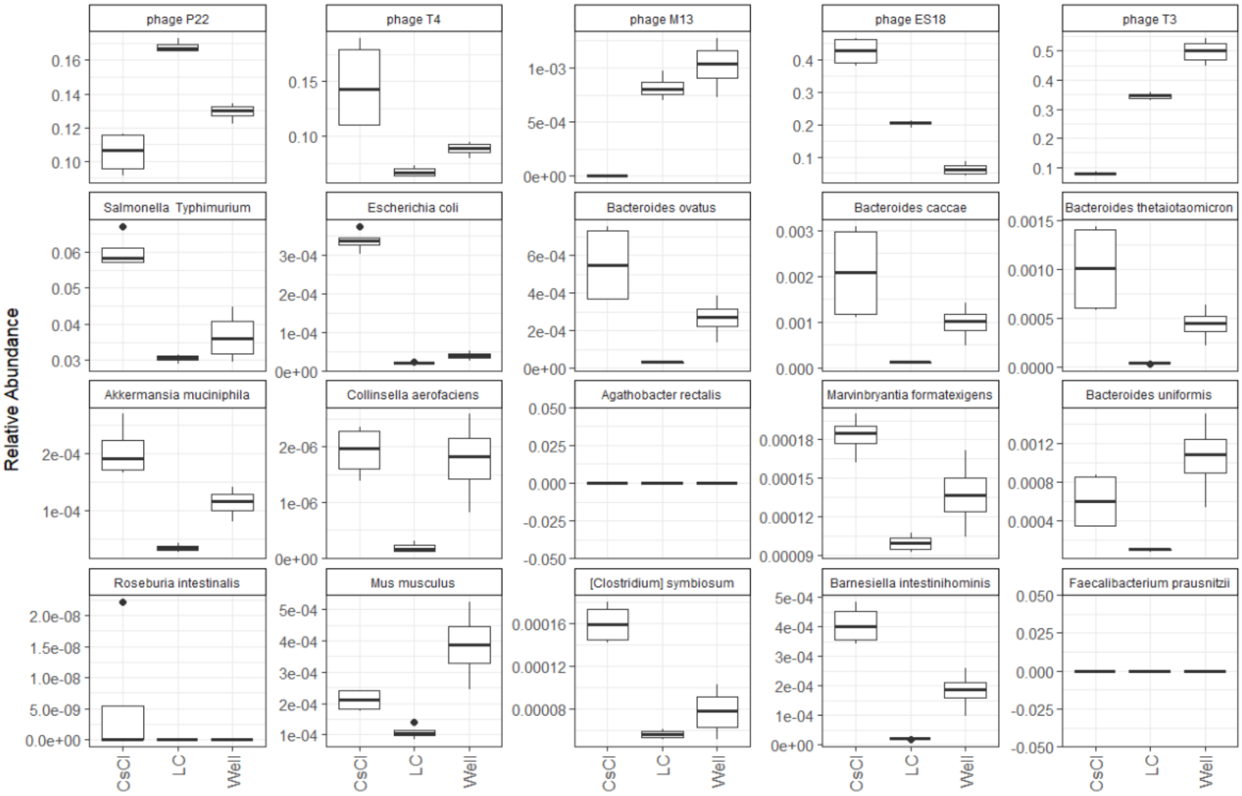


**Supplementary figure 3 - Relative abundance of phage spike-in and bacterial community members in purified VLP-fractions from the gnotobiotic mouse fecal samples.** CsCl density gradient ultracentrifugation (CsCl), anion exchange (AEX) liquid chromatography (LC), and AEX 24-well plates (Well).

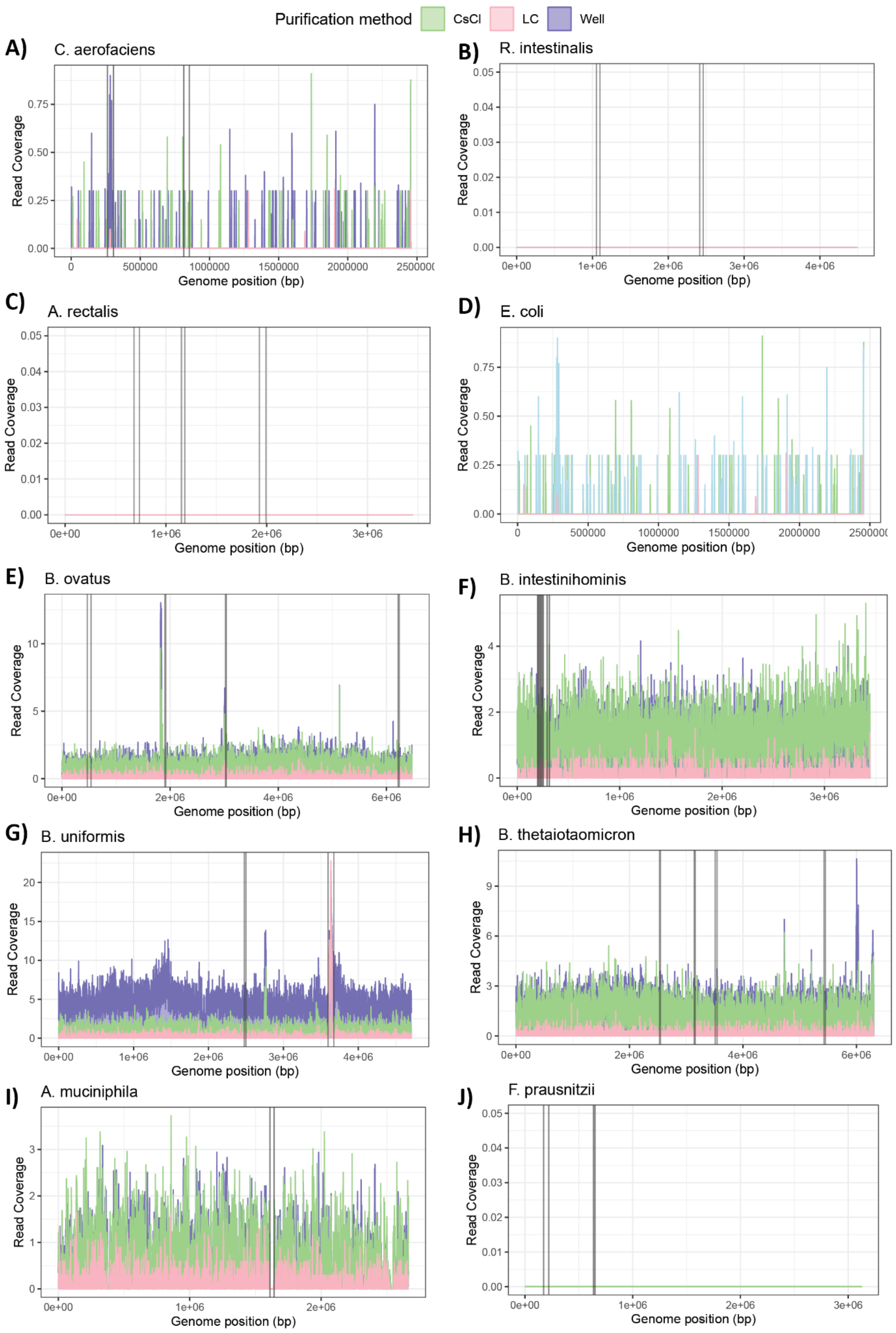
Purification method
CsCl
LC
Well
A)
C. aerofaciens
Read Coverage
0.00
0.25
0.50
0.75
0
500000
1000000
1500000
2000000
Genome position (bp)
B)
R. intestinalis
Read Coverage
0.00
0.01
0.02
0.03
0.04
0.05
0e+00
1e+06
2e+06
3e+06
4e+06
Genome position (bp)
C)
A. rectalis
Read Coverage
0.00
0.01
0.02
0.03
0.04
0.05
0e+00
1e+06
2e+06
3e+06
Genome position (bp)
D)
E. coli
Read Coverage
0.00
0.25
0.50
0.75
0
500000
1000000
1500000
2000000
Genome position (bp)
E)
B. ovatus
Read Coverage
0
5
10
0e+00
2e+06
4e+06
6e+06
Genome position (bp)
F)
B. intestinihominis
Read Coverage
0
2
4
0e+00
1e+06
2e+06
3e+06
Genome position (bp)
G)
B. uniformis
Read Coverage
0
5
10
15
20
0e+00
1e+06
2e+06
3e+06
4e+06
Genome position (bp)
H)
B. thetaiotaomicron
Read Coverage
0
3
6
9
0e+00
2e+06
4e+06
6e+06
Genome position (bp)
I)
A. muciniphila
Read Coverage
0
1
2
3
0e+00
1e+06
2e+06
Genome position (bp)
J)
F. prausnitzii
Read Coverage
0.00
0.01
0.02
0.03
0.04
0.05
0e+00
1e+06
2e+06
3e+06
Genome position (bp)

**Supplementary figure 4 - Genome coverage of bacterial community members in CsCl, LC and Well purified VLP-fractions.** Read coverage for replicate 4 of CsCl density gradient ultracentrifugation (green), anion exchange (AEX) liquid chromatography (LC) (pink) and AEX 24-well plates (blue). Locations of geNomad identified prophage marked with black vertical lines. A) *Collinsella aerofaciens, B) Roseburia intestinalis, C) Agathobacter rectalis, D) Escherichia coli, E) Bacteroides ovatus, F) Barnesiella intestinihominis, G) Bacteroides uniformis, H) Bacteroides thetaiotaomicron, I) Akkermanisa muciniphila*, J) *Facecalibacterium prausnitzii.*

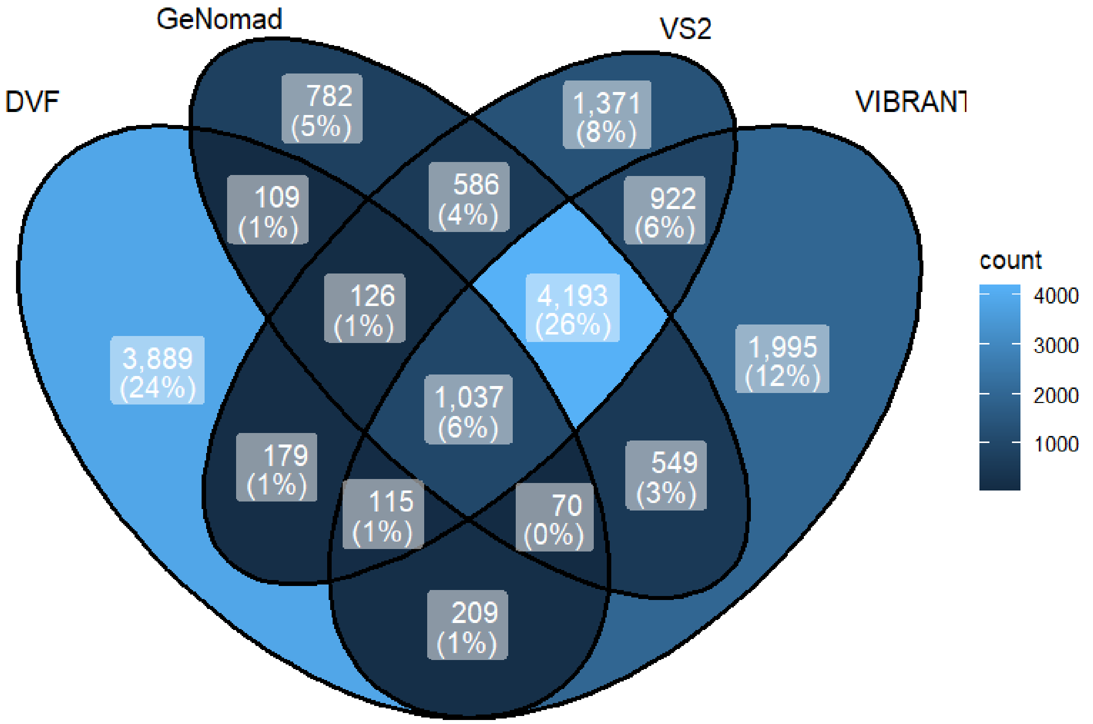


**Supplementary figure 5 - Venn Diagram of LC-purified viral contigs identified with different bioinformatic tools in fecal pellet VLPs from conventional mice.** 16,132 contigs were classified as viral by at least one tool whereas 8,095 were classified as viral by more than two tools. DVF= DeepVirFinder, VS2=VirSorter2

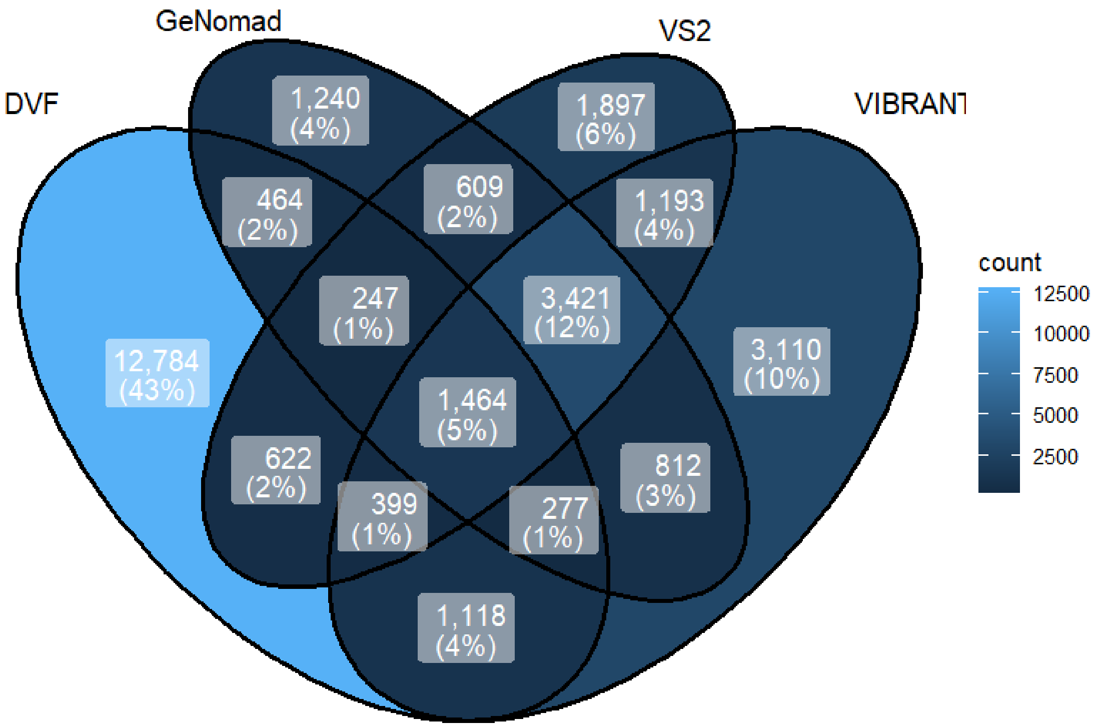


**Supplementary figure 6 - Venn Diagram of CsCl-purified viral contigs identified with different bioinformatic tools in fecal pellet VLPs from conventional mice.** 29,658 contigs were classified as viral by at least one tool and 10,627 were classified as viral by more than two tools. DVF= DeepVirFinder, VS2=VirSorter2

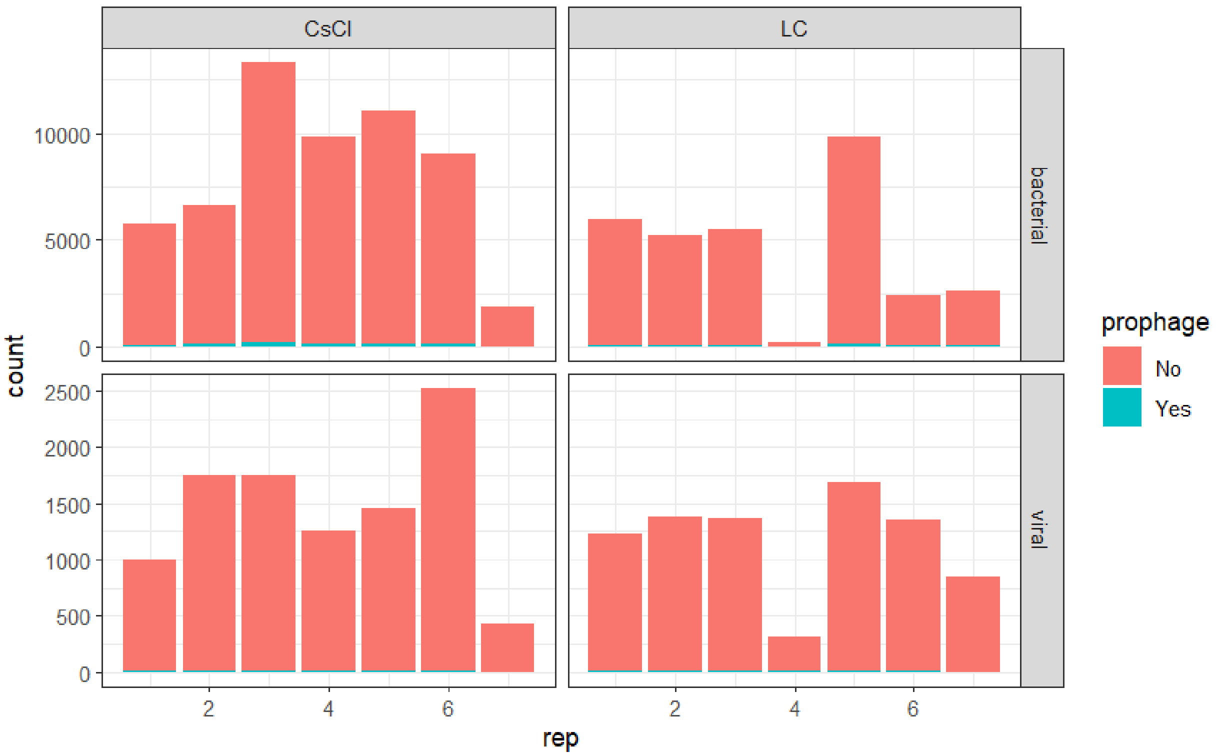


**Supplementary figure 7 - Count of viral and bacterial contigs identified as prophage by CheckV in CsCl and LC assemblies.** Assemblies generated from virus-like particle-fractions purified with either CsCl density gradient ultracentrifugation (CsCl) or anion exchange liquid chromatography (LC)

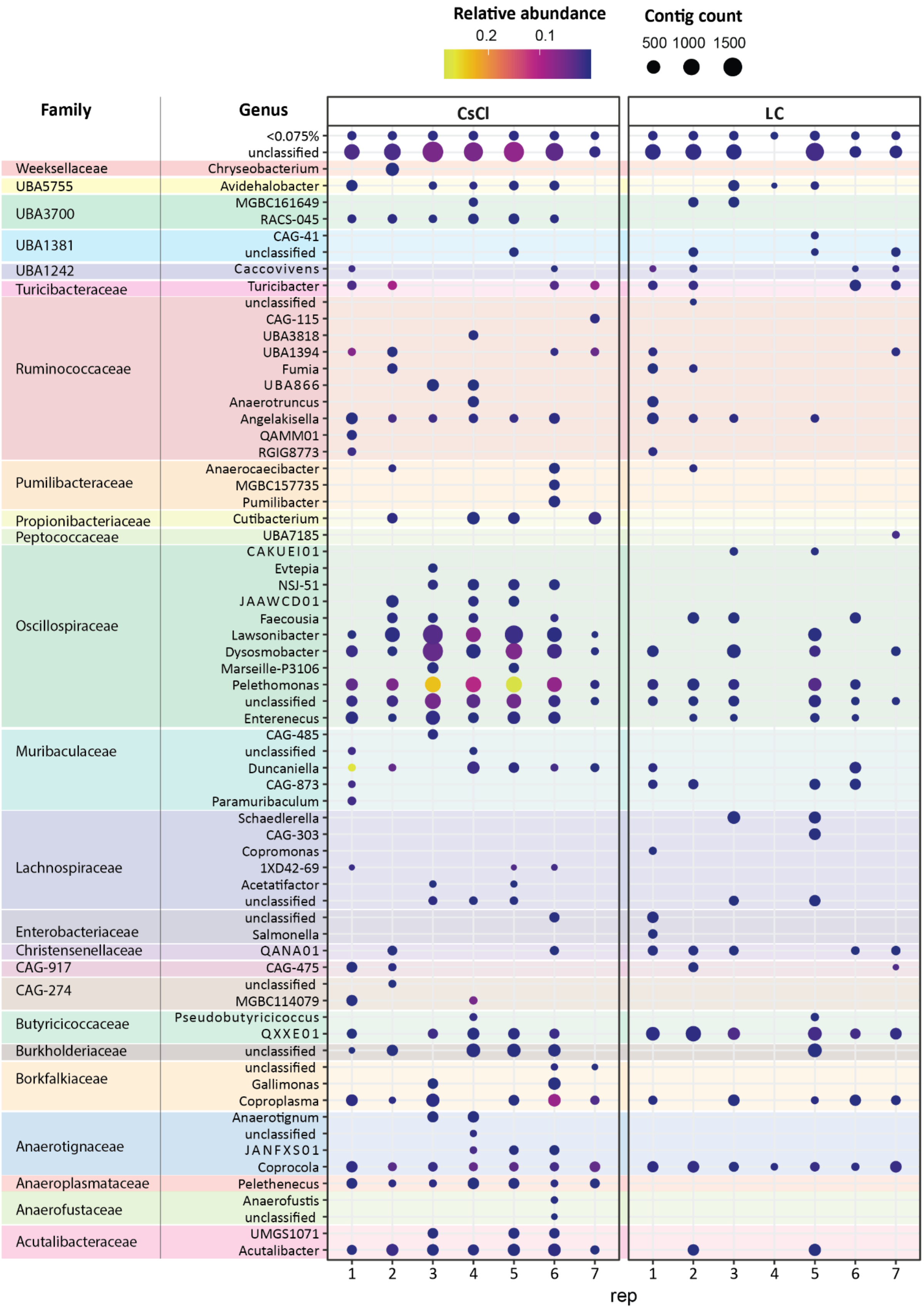
Relative abundance
0.2
0.1
Contig count
500 1000 1500
Family
Genus
CsCl
LC
<0.075%
unclassified
Weeksellaceae
Chryseobacterium
UBA5755
Avidehalobacter
UBA3700
MGBC161649
RACS-045
UBA1381
CAG-41
unclassified
UBA1242
Caccovivens
Turicibacteraceae
Turicibacter
Ruminococcaceae
unclassified
CAG-115
UBA3818
UBA1394
Fumia
UBA866
Anaerotruncus
Angelakisella
QAMM01
RGIG8773
Pumilibacteraceae
Anaerocaecibacter
MGBC157735
Pumilibacter
Propionibacteriaceae
Cutibacterium
Peptococcaceae
UBA7185
Oscillospiraceae
CAKUEI01
Evtepia
NSJ-51
JAAWCD01
Faecousia
Lawsonibacter
Dysosmobacter
Marseille-P3106
Pelethomonas
unclassified
Enterenecus
Muribaculaceae
CAG-485
unclassified
Duncaniella
CAG-873
Paramuribaculum
Lachnospiraceae
Schaedlerella
CAG-303
Copromonas
1XD42-69
Acetatifactor
unclassified
Enterobacteriaceae
unclassified
Salmonella
Christensenellaceae
QANA01
CAG-917
CAG-475
CAG-274
unclassified
MGBC114079
Butyricicoccaceae
Pseudobutyricicoccus
QXXE01
Burkholderiaceae
unclassified
Borkfalkiaceae
unclassified
Gallimonas
Coproplasma
Anaerotignaceae
Anaerotignum
unclassified
JANFXS01
Coprocola
Anaeroplasmataceae
Pelethenecus
Anaerofustaceae
Anaerofustis
unclassified
Acutalibacteraceae
UMGS1071
Acutalibacter
1 2 3 4 5 6 7
1 2 3 4 5 6 7
rep

**Supplementary figure 8 - Specific genera of bacteria present within LC and CsCl-purified VLP-fractions.** CsCl density gradient ultracentrifugation (CsCl) or anion exchange liquid chromatography (LC)

A)

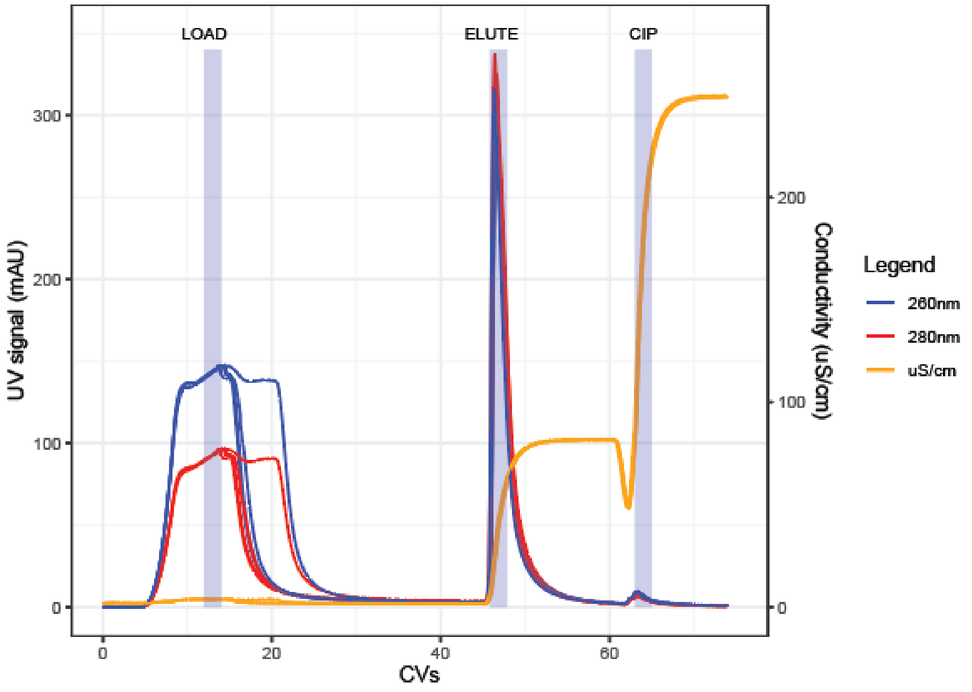


B)

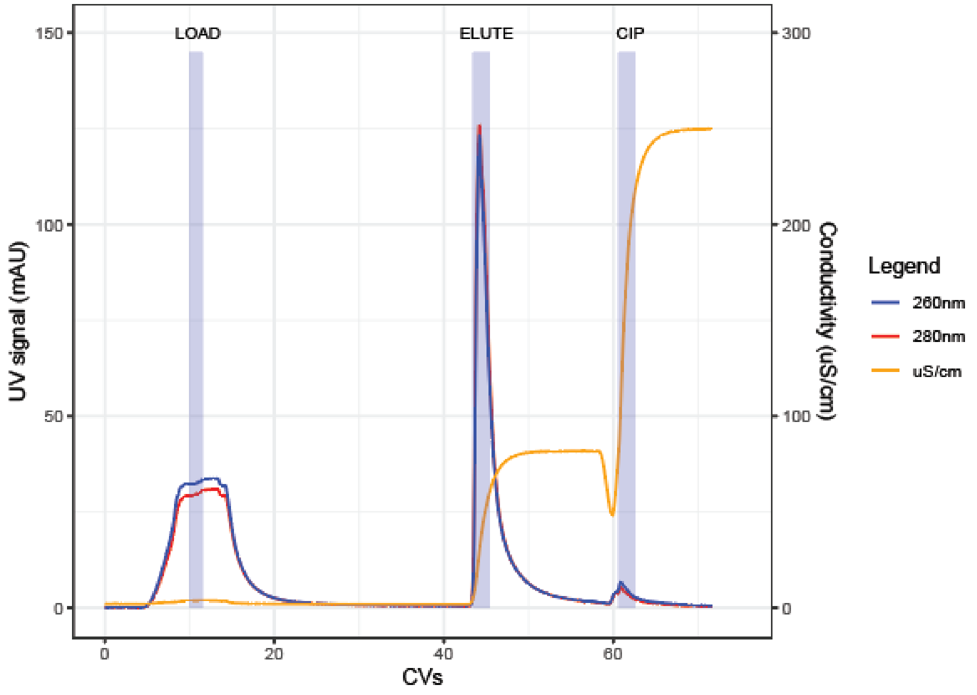

**Supplementary figure 9 - FPLC chromatograms of the simulated microbiome samples containing phage spike-ins.** The collection of samples is indicated with a light blue background. Each collected sample contains 2 ml. A) All 6 phage containing samples (positive) chromatograms are overlaid. B) The negative control chromatogram with no phage-spike in.

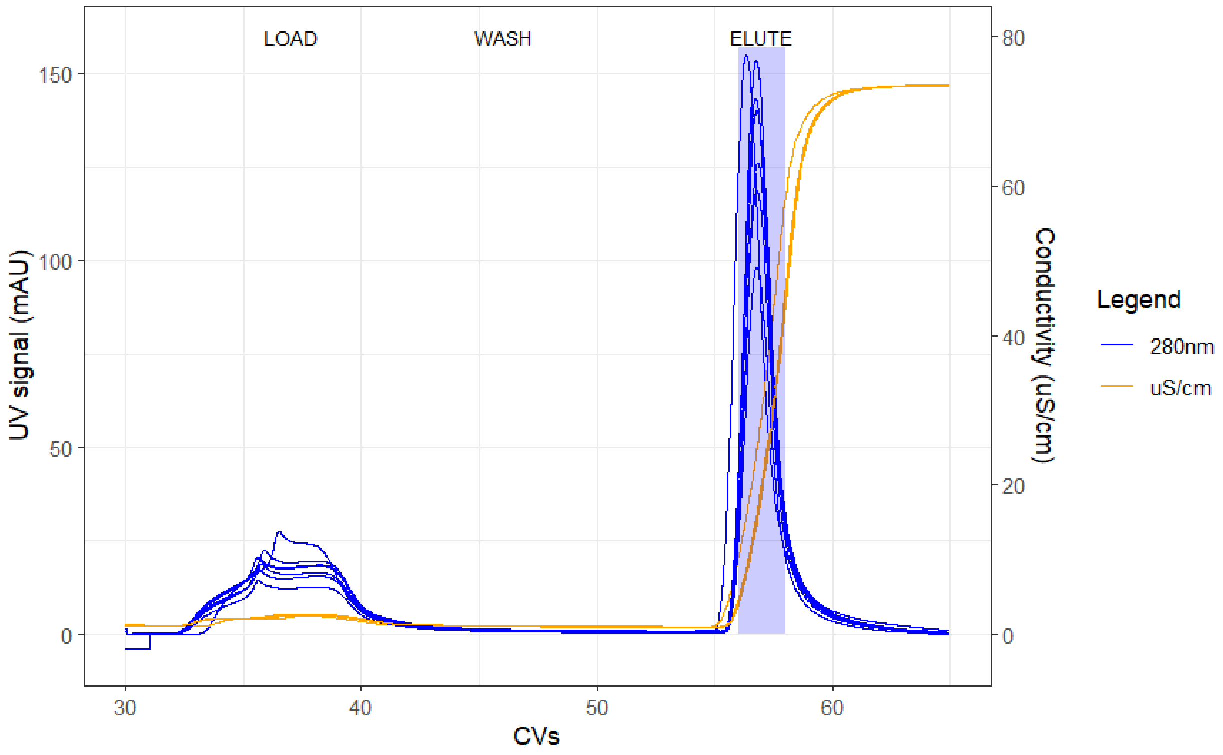


**Supplementary figure 10 - FPLC chromatograms of the conventional murine fecal microbiome samples.** The collection of samples is indicated with a light blue background. Each collected sample contains 2 ml.

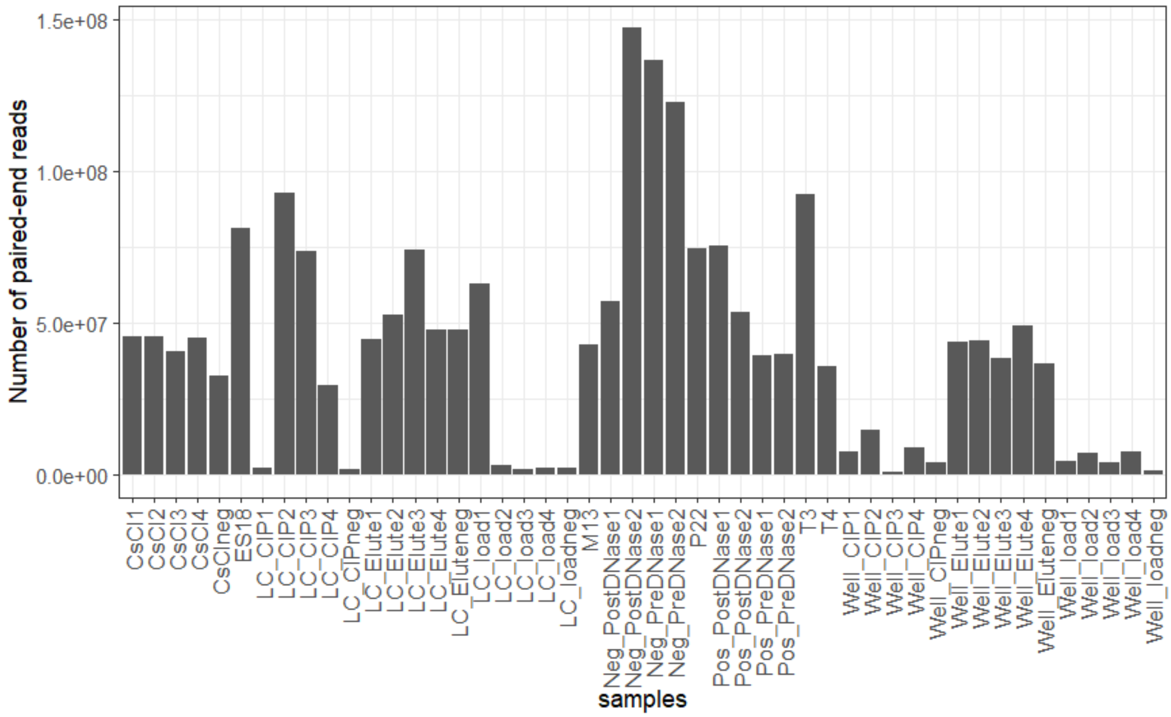


**Supplementary figure 11- Number of trimmed paired-end reads obtained for each of the spike-in samples.** Number indicates replicate number. CsCl = CsCl purified VLPs, LC= anion exchange (AEX) liquid chromatography (LC), Well= AEX 24-well plate. Elute= elute fractions for AEX purified VLPs, Load= Fraction during sample loading of AEX purified VLPs. CIP= Clean-in-place fraction for AEX purified VLPs. Neg= negative control samples with no phage-spike-in. Phage names (P22, ES18, T3, T4, M13) indicate filtered phage lysates.

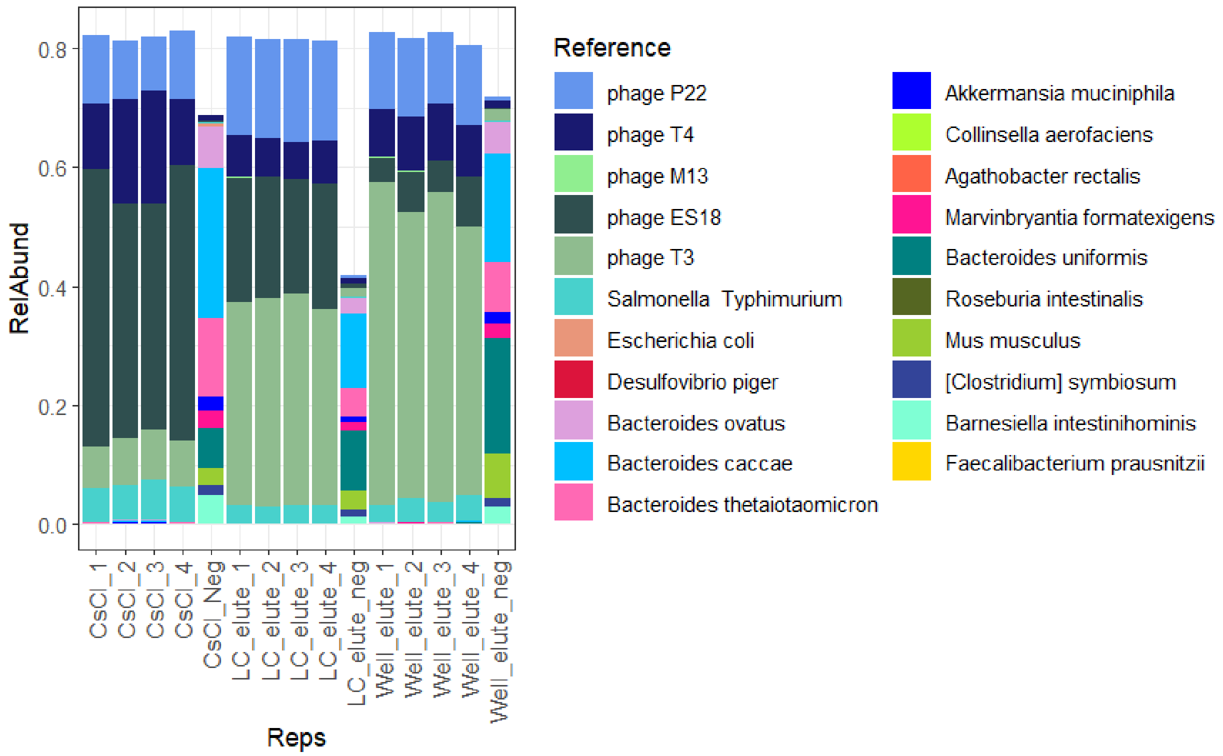


**Supplementary figure 12 - Composition of negative control samples.** Number indicates replicate number. CsCl = CsCl purified VLPs, LC_elute= elute fractions from anion exchange (AEX) liquid chromatography (LC) purified VLPs, Well_elute= elute fractions from AEX 24-Well purified VLPs. Neg= negative control.

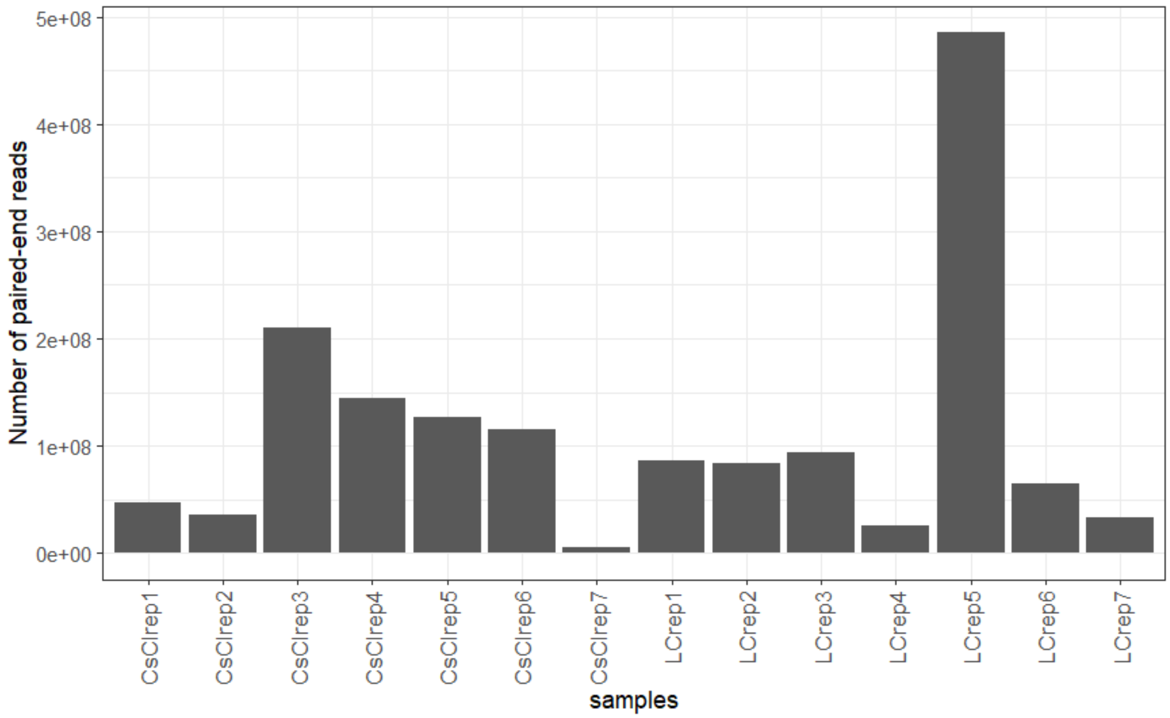


**Supplementary figure 13- Number of trimmed and decontaminated paired-end reads obtained for each conventional murine fecal VLP-fraction sample.** LC= anion exchange liquid chromatography, CsCl= Cesium chloride density gradient ultracentrifugation

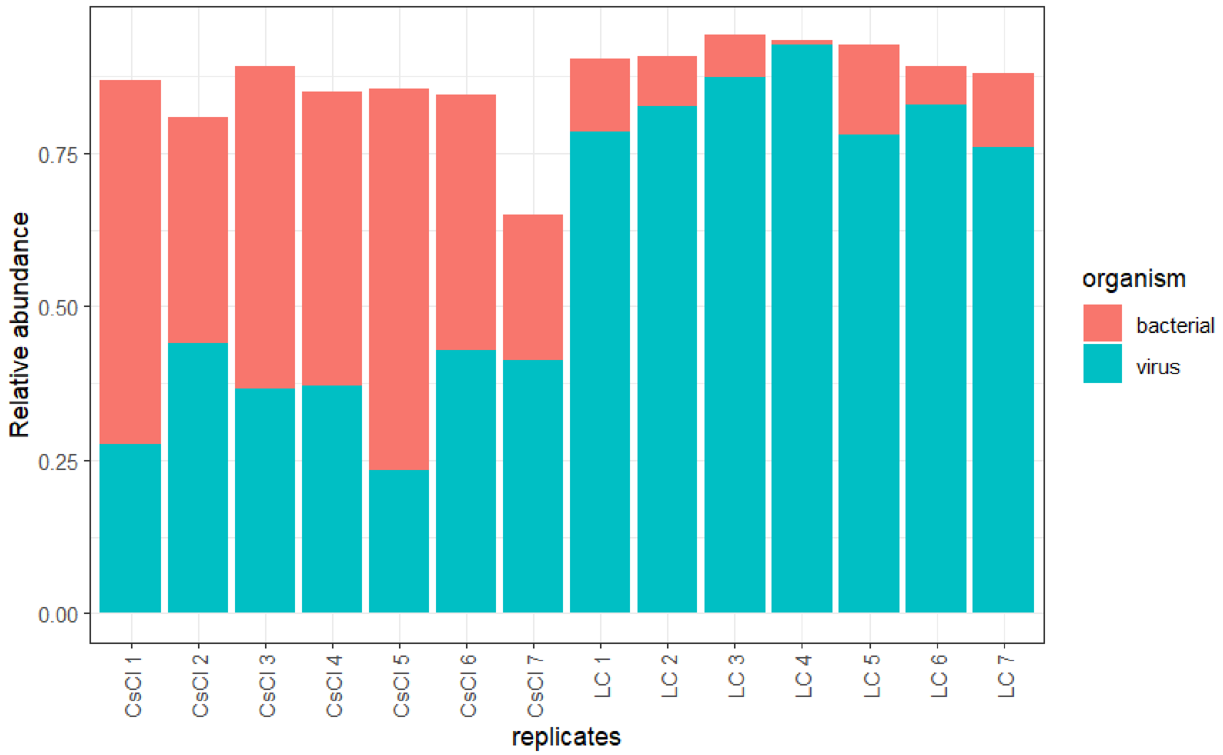

**Supplementary figure 14- Relative abundance of trimmed and decontaminated paired-end reads that map to their associated assemblies for each conventional murine fecal VLP-fraction sample.**

### Supplementary Tables:

**Table S1 - Morphological characteristics of phage used for spike-in**

| Phage | Width (nm) | Length (nm) | Family | Nucleic acid type |
|---|---|---|---|---|
| P22 | 60 | 60 | Podoviridae | dsDNA |
| T3 | 60 | 60 | Podoviridae | dsDNA |
| T4 | 86 | 200 | Myoviridae | dsDNA |
| ES18 | 56 | 266 | Siphoviridae | dsDNA |
| M13 | 6 | 880 | Inoviridae | ssDNA |

**Table S2- The NCBI accession numbers for reference genomes used for phage and bacterial taxa in the gnotobiotic mouse fecal samples**

| Sample | Strain | Reference genome |
|---|---|---|
| M13 phage | NA | GCF_000845205.1 |
| ES18 phage | NA | GCF_000858165.1 |
| P22 phage | NA | GCA_002599445.1 |
| T3 phage | NA | GCF_002745435.1 |
| T4 phage | NA | GCF_000836945.1 |
| *Bacteroides ovatus* | 1896, Type Strain | GCF_001314995.1 |
| *Bacteroides uniformis* | 8492 | GCF_044361425.1 |
| *Bacteroides thetaiotaomicron* | 2079 | GCF_014131755.1 |
| *Bacteroides caccae* | 19024, ATCC43185 | GCF_002222615.2 |
| *Barnesiella intestinihominis* | YIT11860 (JCM15079) | GCF_000296465.1 |

| | | |
|---|---|---|
| *Roseburia intestinalis* | 14610, Type strain L1-82 | GCF_900537995.1 |
| *Eubacterium rectale*<br><br>*(Agathobacter rectalis)* | 17629, A1-86 | GCF_022453685.1 |
| *Faecalibacterium prausnitzii* | 17677, A2-165 | GCF_000154385.1 |
| *Marvinbryantia formatexigens* | 14469, Type strain I-52 | GCF_025148285.1 |
| *Collinsella aerofaciens* | 3979, Type strain | GCF_010509075.1 |
| *Clostridium symbiosum* | 934, Designation 2, ATCC14940 | GCF_000466485.1 |
| *Desulfovibrio piger* | 29098 | GCA_900116045.1 |
| *Escherichia coli* | C | GCF_002079225.1 |
| *Akkermansia muciniphila* | 22959 | GCF_008000975.1 |
| *Mus musculus* | C57BL/6J | GCF_000001635.27 |
| *Salmonella enterica* Typhimurium | LT2 | NC_003197.2 |
